# Characterizing highly dynamic conformational states: the transcription bubble in RNAP-promoter open complex as an example

**DOI:** 10.1101/188524

**Authors:** Eitan Lerner, Antonino Ingargiola, Shimon Weiss

**Affiliations:** Department of Chemistry & Biochemistry, University of California, Los Angeles, CA, USA.; Molecular Biology Institute, University of California, Los Angeles, CA, USA.; Department of Physiology, University of California, Los Angeles, CA, USA.

## Abstract

Bio-macromolecules carry out complicated functions through structural changes. To understand their mechanism of action, the structure of each step has to be characterized. While classical structural biology techniques allow the characterization of a few ‘structural snapshots’ along the enzymatic cycle (usually of stable conformations), they do not cover all (and often fast interconverting) structures in the ensemble, where each may play an important functional role. Recently, several groups have demonstrated that structures of different conformations in solution could be solved by measuring multiple distances between different pairs of residues using single-molecule Förster resonance energy transfer (smFRET) and using them as constrains for hybrid/integrative structural modeling. However, this approach is limited in cases where the conformational dynamics is faster than the technique’s temporal resolution. In this study, we combine existing tools that elucidate sub-millisecond conformational dynamics together with hybrid/integrative structural modeling to study the conformational states of the transcription bubble in the bacterial RNA polymerase (RNAP)-promoter open complex (RPo). We measured microsecond alternating laser excitation μsALEX)-smFRET of differently labeled lacCONS promoter dsDNA constructs. We used a combination of burst variance analysis (BVA), photon-by-photon hidden Markov modelling (H2MM) and the FRET-restrained positioning and screening (FPS) approach to identify two conformational states for RPo. The experimentally-derived distances of one conformational state match the known crystal structure of bacterial RPo. The experimentally-derived distances of the other conformational state have characteristics of a scrunched RPo. These findings support the hypothesis that sub-millisecond dynamics in the transcription bubble are responsible for transcription start site selection.

## I. INTRODUCTION

Biological macromolecules function through changes in their structures. To elucidate their mechanism of action, the structures at all functional steps need to be determined. Atomistic structures are usually solved using X-ray crystallograph^1, 2^or by nuclear magnetic resonance (NMR) spectroscopy^1^. Single-particle cryo-electron microscopy (cryo-EM) has recently been added to this toolkit for characterization and determination of multiple conformations of macromolecules in the ensemble with spatial resolution approaching the atomic leve^13, 4.^Molecular mechanisms could then be inferred from structural “snapshots” attained by these methods (e.g. ^5^). However, although structural snapshots can identify different conformational states (e.g. ligand-bound or unbound, folded or unfolded), they may represent just a subset of all the conformations important for biological function and also lack information on the transitions (and their time scales) between all conformational states.

Biophysical techniques such as NMR, electron paramagnetic resonance (EPR)^6^ and double electron-electron resonance (DEER)^7^ spectroscopies as well as fluorescence-based techniques such as fluorescence polarization^8^, ensemble Förster resonance energy transfer (FRET)^9^, or electron transfer^10^, can provide dynamic information on macromolecular conformations and the dynamics associated with the transitions between them. However, interpreting experimental results derived from these techniques is highly model-dependent and yields limited insight on structural and mechanistic details^11, 12.^ This is mainly because conformational changes in an ensemble are not synchronized, which yields averaged-out signals.

The lack of synchronicity is removed if one molecule is examined at each given time, as afforded by the single-molecule (sm) FRET (smFRET) technique^13, 14^which allows the retrieval of underlying heterogeneity and conformational dynamics. In smFRET, the efficiency of energy transfer from a donor dye to an acceptor dye is measured for each molecule, reporting on the distance between the dyes. Upon attachment of the dyes to different surface residues of the macromolecule, smFRET can report on conformational changes through a distance reaction coordinate between the dyes’ attachment points. However, each pair of dyes accounts for only *one* distance between the attachement points (on the surface of the three-dimensional, 3D, macromolecular structure). In order to capture the full 3D structure and its conformations, multiple such one-dimensional (1D) projections need to be recorded. Indeed, several groups used this approach in conjunction with 3D structures attained from molecular simualtions (e.g. molecular dynamics, Monte Carlo)^15–19^. This approach allows identification and solving of 3D macromolecular structures that are ‘too dynamic’ and have not been characterized by classical structural biology techniques.

If transition rates between different conformatios are slower than the smFRET temporal resolution, they could be resolved. If, however, the conformational states are separated by a low activation barrier, yielding transition rates that are faster than the method’s temporal resolution, they would be averaged-out and become indistinguishable. In smFRET of freely diffusing molecules, fluorescence bursts (of donor and acceptor photons) are generated when the molecules traverse the observation volume. These bursts typically last a few milliseconds. FRET values are calculated for each burst (molecule) with this temporal resolution. Single bursts (molecules) with different FRET efficiencies are then grouped into different subpopulations of distinct conformational states that interconvert at time scales slower than ~10 ms. Faster conformational transitions (occuring at timescales of ~ 0.1-10 ms) are averaged out^20, 21.^Several techniques based on analysis of photon statistics were developed to study conformational dynamics occuring within this faster temporal range^22–24^. Of particular interest to the work presented here, we note the burst variance analysis (BVA) techique which is based on the analysis of the variance of FRET efficiency in within bursts^24^, as well as the photon-by-photon hidden Markov modelling approach (H2MM)^25^ that quantifies conformational dynamics parameters (number of conformational states involved, their FRET efficiencies, and their interconversion rate, down to ~1-10 μs^25^). To characterize the structures of different conformational states in the ensemble, it is therefore important to first identify each conformational state with the highest available temporal resolution, and to do so for multiple differently-labeled constructs.

In this study, we apply this approach to study the conformational dynamics of the transcription bubble in RNAP-promoter open complex (RPo). To initiate DNA transcription starting from a gene’s promoter sequence, RNAP has to first specifically bind to the promoter and then open a transcription bubble by melting a segment of 10-12 base pairs (bp) to form the RPo^26, 27.^While doing so, RNAP positions a pair of bases from the downstream fork of the bubble in front of its active site to dictate the initial sequence of the RNA to be transcribed^28^. Therefore, the size of the transcription bubble imposes which pair of DNA bases will be in front of the active site, hence dictating the (canonical) transcription start site (TSS). We note, however, that in the presence of specific di-nucleotides, TSS can be reprogramed^29–31^. Recently, Robb *et al.* have identified bubble size fluctuations in RPo and suggested that they are involved in TSS selection^32^. They performed smFRET measurements on doubly-labeled promoter dsDNA constructs sensitive of changes in the bubble. BVA analysis has indicated sub-millisecond bubble conformational dymanics, implying that DNA bases situated downstream to the transcription bubble may dynamically melt and transiently be reeled into the active site. Such hypothesized melting and reeling-in mechanism could increase the overall bubble size and is reminescent of ‘DNA scrunching’^33, 34^, but does not require the presence of NTPs. If validated, this mechanism could explain how transcription initiates from positions other than the canonical TSS along the promoter sequence.

Robb *et al.* have also presented results of a gel-based transcription assay for the lacCONS promoter^35^, exhibiting two discrete bands (i.e. transcripts of two different distinct lengths). If these bands correlate with the transcription bubble dynamics, they may imply the existence of two conformational states, one of which is RPo, and the other is a modified RPo having a few additional DNA bases scrunched into the active site. To prove or refute this hypothesis, it is important to quantify the transcription bubble dynamics using a conformational state model and to do so for more than two positions of dyes.

We therefore performed diffusion-based smFRET measurements on a series of such donor-acceptor labeled lacCONS promoter constructs and characterized the donor-acceptor distances for each construct at each conformational state, as well as the interconversion rate constants, by employing the H2MM approach. We then used the attained distances as constraints to test which of the two identified conformational states matches the known RPo crystal structure^36^ and what is the conformation of the other state. Our results support the two-state hypothesis for TSS selection in the lacCONS promoter.

## II. EXPERIMENTAL

### A. μsALEX-smFRET distance measurements

We performed microsecond alternating laser excitation (μsALEX) smFRET measurements^37–39^ on a library of dsDNA constructs having the lacCONS promoter sequence^35^. All of the constructs had the template strand (T) labeled with the acceptor (A) ATTO647N attached to a specific DNA base (denoted here as TA) and the nontemplate strand (NT) labeled with the donor (D) Cy3B at another specific DNA base (denoted here as NTD). The identity of the labeled DNA bases is denoted relative to the promoter canonical TSS, where negative or positive values define regions that are upstream or downstream of TSS respectively. Throughout this work, we therefore use the following nomenclature to name a D-A labeled lacCONS construct: (-3)TA-(-8)NTD is one construct in the library with an A dye labeling the base at register (-3) upstream to the canonical TSS in the template strand and with a D dye labeling the base at register (-8) upstream to the canonical TSS in the nontemplate strand. Table S1 of the supplementary material lists all constructs used for this study.

Measurements of the labeled dsDNA constructs were taken in the absence and in the presence of *Escherichia coli* (*E. coli*) RNAP in order to track conformational changes in the promoter induced by the formation of RPo. As a control, measurements were also taken in the presence of RNAP after incubation with nucleotides (NTPs) to assess promoter escape activity (FIG. S4 of the supplementary material). All measurements were taken at T=25°C, for ~30 minutes, and with the labeled dsDNA at a concentration of ≤100 pM. Additional preparative and experimental details can be found elsewhere^40, 41.^

D-A distances (*r̅_D-A_*) for all constructs in RPo (and for free promoter as controls) were extracted according to the steps outlined in supplementary material.

### B. FRET-restrained positioning and screening

In this study, we measured multiple distances for dsDNA promoter in the presence of the bacterial RNAP in the RPo state, for which a crystal structure already exists (pdb code: 4XLN)^36^. After calculating *r̅_D−A,E_* values for each conformational state (as identified by H2MM and transformed to *r̅_D−A,E_* using the Eqs. S16-S18, S21) for all D-A labeled constructs, distances were classified into different groups, where each group presumably defines one conformational state. As can be seen in (FIG. 3 and Table S2 of the supplementary material) two groups were identified as RNAP-bound conformations. Since our results indicate that the transcription bubble has two conformations, we wanted to identify which of the smFRET - derived conformational states represents the solved structure (as represented in pdb code: 4XLN^36^) and to what extent and how the other conformation deviates from it.

As already demonstrated, a group of *r̅_D−A_* distances can be used as constraints for structural determination by computation methods such as molecular dynamics (MD)^17, 42–47,^ Monte Carlo (MC)^18, 48^ and coarse-grained (CG)^49–51^ simulations as well as using structural databases^52^ or distance-constrained triangulation^53^. *r̅_D−A_* reports on mean distance between the D and A ***fluorophore centers***, rather than the mean distance between ***DNA bases***, an additional computational step that takes into account the dye linker length and all possible dye configurations in space (dye accessible volume, or AV) is needed for this ‘distance translation’. Such a tool, dubbed “the FRET-restrained positioning and screening (FPS) approach” was developed by Kalinin et al.^17^ and adapted for this study.

The FPS software calculates the AV of a dye given the atom to which it is attached to in a given pdb structure, the simplified geometrical (ellipsoid) parameters of the dye, and its linker length *L_link_* and width *w_link_*. In our case, *L_link_* = 20 Å and *w_link_* = 4.5 Å for Cy3B while *L_link_* = 20.5 Å and *w_link_* = 4.5 Å for ATTO647N. The ellipsoid parameters for Cy3B are {R_;_ = 6.8 Å; *R_2_* = 3.0 Å; *R_3_* = 1.5 Å } and for ATTO647N {R_;_ = 7.15 Å; *R_2_* = 4.5 Å; *R_3_* = 1.5 Å }. These parameters are taken from the table provided in the FPS software manual (for Cy3B, we used the Cy3 values).

The AV surface provides a set of possible positions for the fluorophore to assume in space. Each pair of such positions for D and A, makes up one possible *r*_D−A_. However, it is expected that D and A will explore all possible orientations within their AVs, at times much faster (nanoseconds) than the best time resolution H2MM can offer (~10 μs, the time between consecutive photons in our case). Therefore, the expected value *r̅_D−A_* (and the expected standard deviation, *SD*(*r̅_D−A_*)) are calculated by averaging over all possible pairs of dye positions in the context of their AVs. These averages are then compared to expected distances from the known structure.

We compared the set of apparent mean D-A distances, *r̅_D−A,E_*, values that were derived from experimental mean FRET efficiencies, *Ē* (Table S2 of the supplementary material), for each conformational state, with the mean distances *r̅_D−A_,* that is expected between these dyes based on bacterial RPo structure^36^ (see Eqs. S17-S20 and discussion in the supplementary material). See Eqs. S17, S18 & S21, FIG. S17 and discussion therein on the validity of the *r̅_D−A,E_* approximation. The crystal structure includes the promoter T and NT strands and all the subunits that make up the RNAP holoenzyme. We separated the crystal structure into T, NT and RNAP holoenzyme, as three distinct entities. Then, using the FPS software, we tried to reconstruct the complex using the experimentally-derived distance constraints while minimizing the RMSD values between all experimental and expected D-A distances.

It is worth noting that the FPS software uses two measures of distances interchangeably, namely the mean D-A distance, *r̅_D−A_,* and *r_mp_*, the distance between the mean positions of the dyes. The two measures deviate from each other, especially at short distances and the FPS software uses a 3^rd^ order polynomial approximation to allow conversion between the two^17^. In this work we use the *r̅_D−A_* distance measure.

## III. RESULTS AND DISCUSSION

### A. The screening process

We measured a library of 43 dsDNA constructs doubly-labeled (D-A) at different positions of the lacCONS promoter^35^. We used all 43 constructs for μsALEX-smFRET measurements of free promoter, calibrating the correction factors *lk, dir* and γ (see Table S1 and *μsALEX-smFRET* in the supplementary material). However, out of these combinations, only 23 constructs had D and A dyes’ positions that are expected (based on the crystal structure) to exhibit different *r̅_D−A_* in RPo as compared to the free DNA (colored rows in Table S1 of the supplementary material). The constructs included D-A pairs positioned along two different reaction coordinates: (i) ***scrunching coordinate***, where one dye is attached to a DNA base (T or NT) upstream of the transcription bubble or at the edge of the bubble (promoter registers ≤ -8) and the other is attached to a DNA base (T or NT) downstream of the transcription bubble or at the edge of the bubble (promoter registers ≥ -2; FIG. 1a and Table S1 of the supplementary material); (ii) ***bubble coordinate***, where both dyes are attached to T and NT on registers inside the transcription bubble in RPo (promoter registers -10 to +1; FIG. 1b and Table S1 of the supplementary material); Bubble coordinate constructs in RPo are expected to show an increase in *r̅_D−A_* due to bubble opening, while scrunching coordinate constructs in RPo are expected to show a decrease in *r̅_D−A_* due to reeling of DNA into the enzyme (as compared to free DNA). However, in some cases, degeneracies could be expected where *r̅_D−A_* values in free promoter and in RPo are similar (FIG. 3, black filled circles vs. red filled circles).

**FIG. 1:**
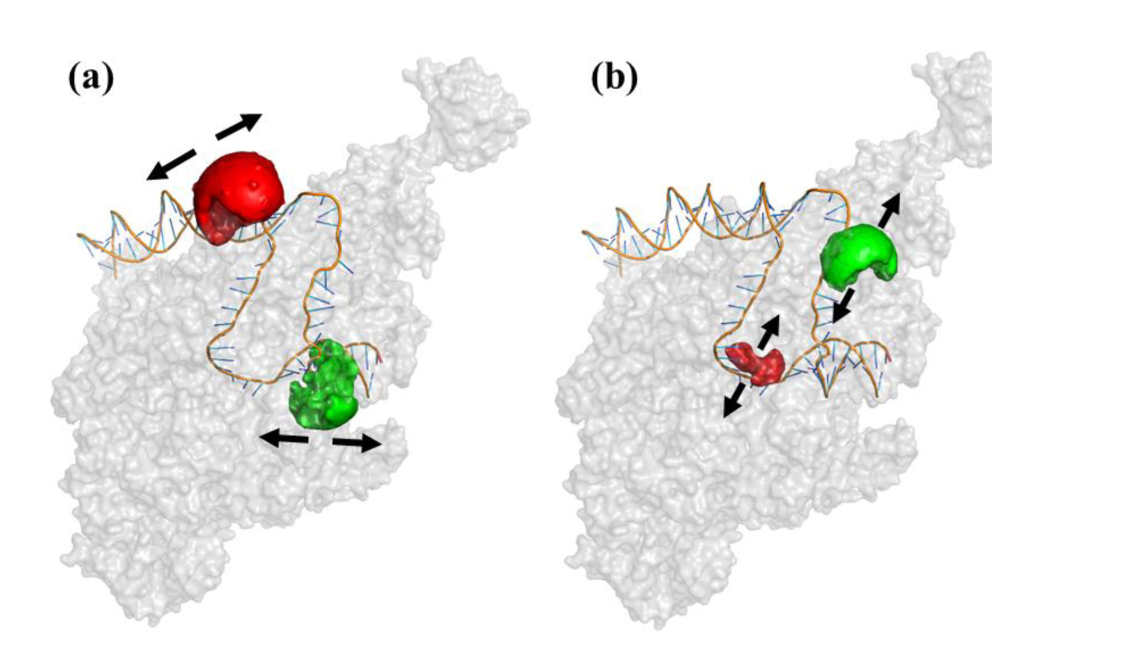
D-A labeled promoter constructs representing two reaction coordinates. Examples of dyes’ attachment points in Po for two reaction coordinates (pdb code: 4XLN). (a) scrunching coordinates: dyes attached to bases upstream and downstream relative to the transcription bubble; (b) bubble coordinates: dyes attached to bases within the transcription bubble. The arrows show the general directions in which distance changes are expected in the two reaction coordinates. Dye AVs are also shown as green (D) and red (A) partial spheres.

These constructs are expected to be sensitive to dynamic distance changes due to possible fluctuations in the bubble size or fluctuations in scrunching (FIG. 1). We note, however, that some of the constructs could be insensitive to these fluctuations (due to FRET limited dynamic range).

We further screened the 23 constructs according to the following criteria: (i) the *PR* histogram of free promoter yields a single Gaussian population (except for cases in which quenching of FRET occurs^54^; *PR* histograms of free promoter in FIGs. 2a and S2). FIG. S3 provides an example of such a dsDNA construct; (ii) <u>bubble opening activity</u>: *PR* histograms of RPo yield two subpopulations, one similar to that of the free promoter and the other representing the RNAP-bound fraction, with a peak *PR* value different than that of the free promoter subpopulation (comparing *PR* histograms of free promoter and RPo in FIGs. 2, S2 and S4 of the supplementary material). FIG. S5 of the supplementary material lists all constructs that did not show sufficient bubble opening activity; (iii) qualitative test for conformational dynamics: a majority of sm bursts in the *PR* subpopulation representing the RNAP-bound fraction exhibit *σ(PR)* values larger than expected from shot noise (burst variance analysis, BVA, plots of RPo in FIGs. 2 and S2; more information on BVA in supplementary material); constructs that exhibited a signature of dynamics in BVA plots in free promoter form (FIG. S6 of the supplementary material), or those that did not exhibit any signature of dynamics in BVA plots in the RNAP-bound *PR* subpopulation (FIG. S7 of the supplementary material), were not selected; (iv) promoter escape activity: the RPo *PR* subpopulation is significantly decreased after addition of NTPs, and the resulting distribution resembles the *PR* of the free promoter (comparing *PR* histograms of RPo and RPo + NTPs in FIG. S4 of the supplementary material); (v) the experimental results are converted from H2MM-retrieved most likely *PR* values to Ē values (Eq. S16 of the supplementary material) and from it to apparent mean D-A distance, *r̅_D−A,E_* (using the procedure described in Eqs. S17, S18, S21 of the supplementary material), and collectively compared against RPo crystal structure (pdb code: 4XLN)^36^. This structure includes part of the promoter with registers -35 through +12, hence scrunching coordinate constructs with dyes at registers outside this range ((+15)TA-(-15)NTD and (+15)TA-(-8)NTD in our case) are excluded. 9 constructs out of 23 scrunching/bubble coordinate constructs were selected following these screening steps (yellow-shaded rows, Table S1 of the supplementary material). Out of the 9, 6 were bubble coordinate constructs and 3 were scrunching coordinate constructs (FIGs. 2, S2, S4, and yellow-shaded rows of Table S1 of the supplementary material).

In summary: the final 9 selected constructs exhibited a unique subpopulation in the *PR* histograms that represents the RNAP-bound fraction. This subpopulation exhibited dynamics (as assessed by BVA) while the free promoter fraction did not exhibit such dynamics (FIGs. 2 and S2 of the supplementary material). We therefore conclude that these bursts carry information on fast interconverting conformational states and that the number of different conformational states cannot be extracted from simple fits of *PR* histograms to a sum of Gaussians. BVA allowed us to identify the presence of fast dynamics; the next section shows how to extract quantitative dynamics parameters.

### B. Quantification of underlying dynamics using H2MM

Photon-by-photon hidden Markov modelling (H2MM) analysis was applied to smFRET photons bursts of all 9 selected constructs in RPo (for more information on H2MM, see supplementary material). To explain the results of the H2MM analysis we choose to follow the results for (-3)TA-(-6)NTD (bubble coordinate) as an example. In this construct, the free promoter fraction was not quenched and hence showed up as a subpopulation with high *PR* values in the *PR* histograms (FIG. 2a, center column). Additionally, this construct was expected to have a D-A distance in the free promoter that is in within the FRET measurable region, therefore FRET is not quenched in the free promoter form (FIG. 3, black filled circles). As expected, this peak did not exhibit dynamics in the BVA plots (FIG. 2a, center column), and therefore was assigned a single conformational state. Upon addition of RNAP, a subpopulation with lower *PR* values emerged (RPo; FIG. 2b, center column). This subpopulation exhibited dynamics as determined by BVA (FIG. 2b, center column). It was therefore assumed to exhibit at least two conformational states (interconverting with rate constants in the range ~10^4^-10^2^ s^-1^). The model, therefore, should have at least three states (Min. *q,* Table S2 of the supplementary material), with one free promoter state and two (or more) RNAP-bound conformational states.

**FIG. 2:**
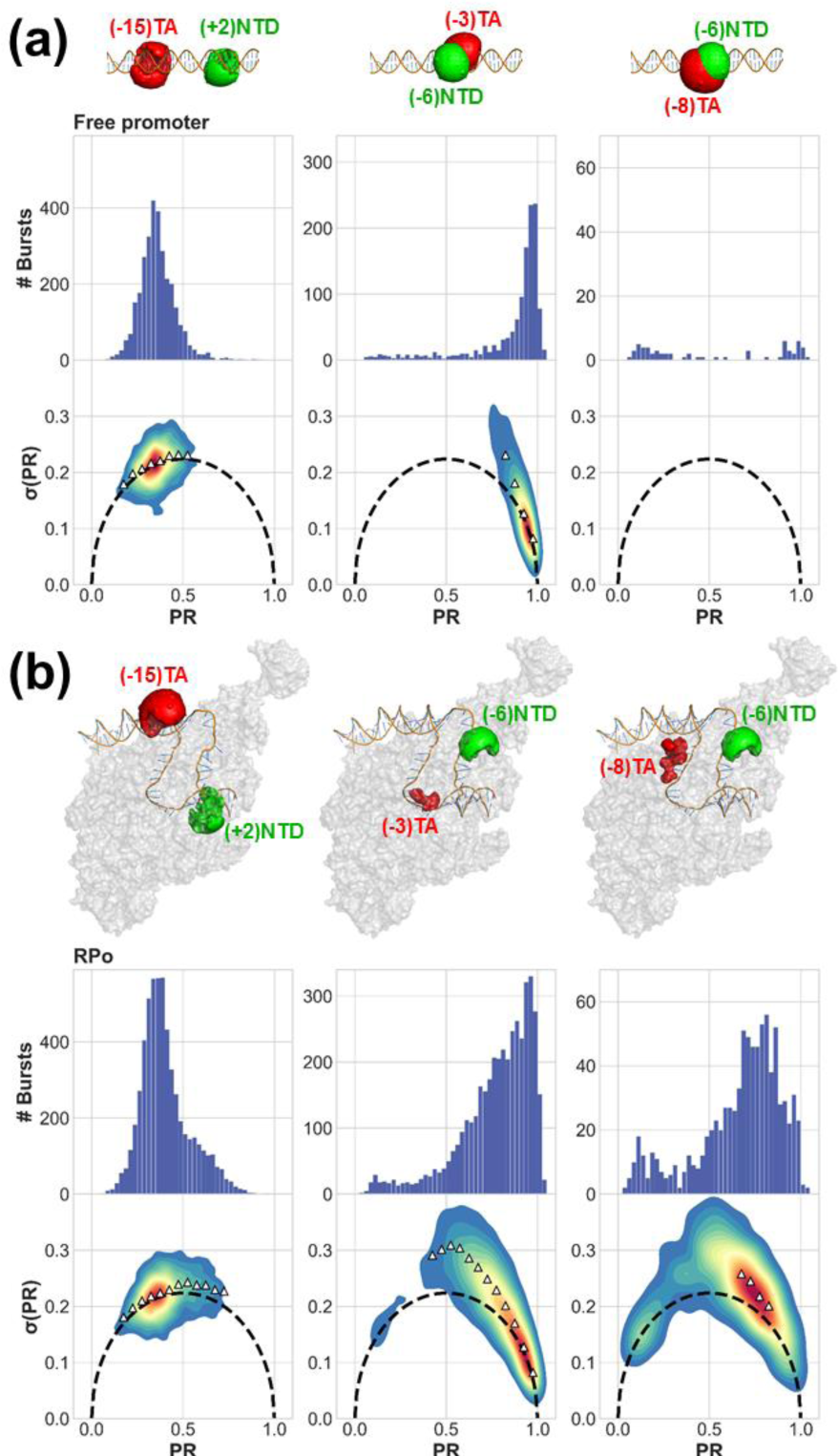
Qualitative assessment of smFRET dynamics using BVA. Proximity ratio (*PR*) histograms and BVA plots of σ(*PR*) versus *PR* show that. (a) free promoter is either characterized by a single *PR* population with no smFRET dynamics or with no *PR* population at all, due to quenching of FRET, and that (b) RNAP bound to promoter DNA induces a unique PR subpopulation that exhibits smFRET dynamics. Depiction of differently labeled D-A constructs in free promoter (a) and RPo (b) states, were generated by calculating D and A AVs (green and red surfaces, respectively) on top of DNA labeling positions in the context of RPo crystal structure (pdb code: 4XLN).

Using H2MM, the most likely values of the *PR* and rate constant values were retrieved for each state using a given state-model. Since this construct represents the bubble coordinate, we expected the most likely *PR* value of the RNAP-bound states to be smaller or equal relative to the value of the free promoter. Using the *BIC’* criterion (Eq. S11) requiring *BIC’<0.005,* the 4 states model was the state-model with the minimal number of parameters that reached a value of *BIC’,* lower than the 0.005 threshold. Therefore, the 4-state model was found to be a better model to describe the underlying conformational dynamics in the construct (-3)TA-(-6)NTD (FIG. S8 and Table S2 of the supplementary material). To validate the use of the *BIC’* statistical criterion for choosing the most appropriate number of states, we performed H2MM analyses of simulated smFRET experiments including dynamics (FIG. S9-S11 and Table S3 of the supplementary material). 8 constructs were best described by a 3-state model and 1 construct was best described by a 4-state model. Results for all constructs are summarized in Table S2 of the supplementary material. The next section outlines an approach for classifying these results into distinct groups of *r̅_D−A,E_* values that represent different conformations.

### C. Classification of D-A distances to conformational states of the transcription bubble

The *r̅_D−A,E_* values (FIG. 3) derived from results of H2MM analyses (states and rate constants derived from H2MM) indicate that: (i) the *PR* subpopulation of RPo (FIGs. 2b and S2, and Table S2 of the supplementary material) is an average of two dynamically interconverting states; (ii) in all constructs, except for (-15)TA-(+2)NTD, one out of the two conformational states, associated with the RNAP-bound subpopulation, had *r̅_D−A,E_* values that coincided with the values expected from the crystal structure of RPo (FIG. 3, magenta open squares vs. red filled circle). We therefore give this state the name *RPo state*; (iii) in all bubble coordinate constructs, except for (-3)TA-(-8)NTD, the second state had larger *r̅_D−A,E_* values as compared to *RPo state r̅_D−A,E_* values (FIG. 3, blue open diamonds vs. magenta open squares); (iv) in all scrunching coordinate constructs, the second state had smaller *r̅_D−A,E_* values as compared to *RPo state r̅_D−A,E_* values (FIG. 3, blue open diamonds vs. magenta open squares). When taken together (except for (-3)TA-(-8)NTD), these results suggest that the second state resembles DNA scrunching: distances of scrunching coordinates decrease while distances of bubble coordinates increase (as compared to the RPo state, FIG. 3, blue open diamonds). We therefore name this state *scrunched RPo state.* The bubble-coordinate construct (-3)TA-(-8)NTD exhibited a decrease in distance relative to RPo state. This exception may hint to the fine details of topological changes inside (or outside of) the transcription bubble in the *scrunched RPo state.*

**FIG. 3:**
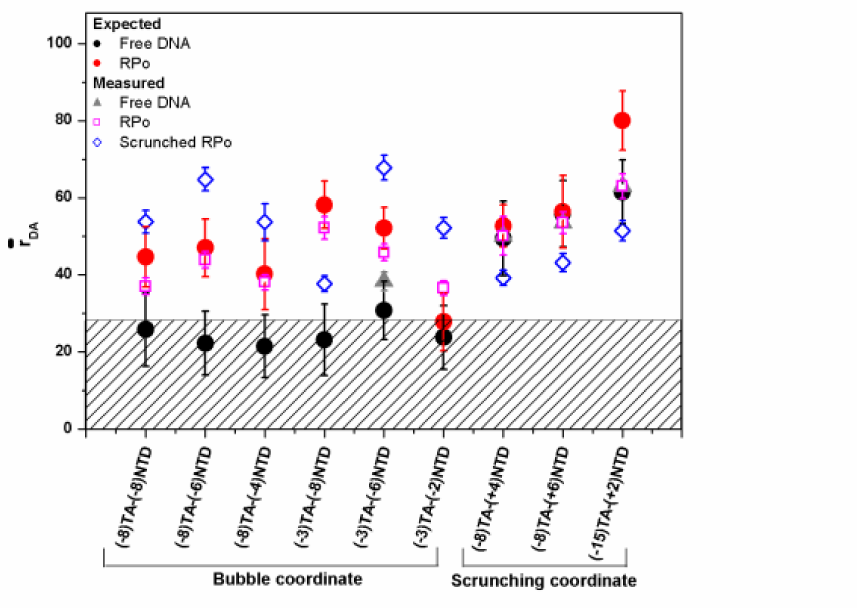
Comparison of measured apparent D-A distances, *r̅_D−A, E_* with expected D-A distances *r̅_D−A_*, The experimentally-derived *r̅_D−A, E_* values of different conformational states for each D-A labeled lacCONS constructs are compared with the expected D-A distances from RPo crystal structure (pdb code: 4XLN) in free DNA (black dot) and in RPo (red dot). Two conformational states were found (with sub-millisecond underlying interconversion dynamics) in the RNAP-bound fraction. Their derived distance values are shown (magenta open squares) for the subset that defines a conformation closest to the known RPo structure. Distances for the other conformation are also shown (blue open diamonds). The values of the free promoter fraction are also presented (grey triangles; for the D-A labeled promoter constructs that do not exhibit quenched FRET). Quenched FRET constructs had an expected D-A distance below 0.5R_0_ (shown as shaded area). The error bars in the measured distances represent experimental standard error (Eq. S21). The error bars in the expected distances represent the D-A standard deviation, as calculated from all possible donor and acceptor positions in space (dye AVs).

Next, the two sets of *r̅_D−A, E_* values that were assigned to the *RPo state* and to the *scrunched RPo state* were used as distance constraints in FRET-restrained positioning and screening (FPS)^17^, as described below.

### D. Assessment of FRET-derived conformational states against a known RPo structure

In principle, the FRET-derived *r̅_D−A, E_* values that were assigned to the *RPo state* should agree with *r̅_D−A, E_* values of the crystal structure of bacterial RPo^36^. Indeed, and as mentioned above, this assertion seems to be true except for (-15)TA-(+2)NTD (FIG. 3, magenta open squares vs. red filled circles). A more quantitative assessment was performed using the FPS approach (see supplementary material) to verify which of the conformational states’ *r̅_D−A, E_* sets *(RPo state* and the *scrunched RPo state*) better agrees with the known RPo crystal structure. FPS fitting results demonstrated that distances of the *RPo state* afforded reconstruction of the promoter DNA conformation that coincided with the known crystal structure with only minimal deviations (RMSD=4.2 Å; FIG. 4a), hence justifying the assignment. FPS fitting results demonstrated that distances of the *scrunched RPo state* afforded reconstruction of the promoter DNA conformation that had large deviations from the known crystal structure (RMSD=14.0 Å; FIG. 4b).

**FIG. 4:**
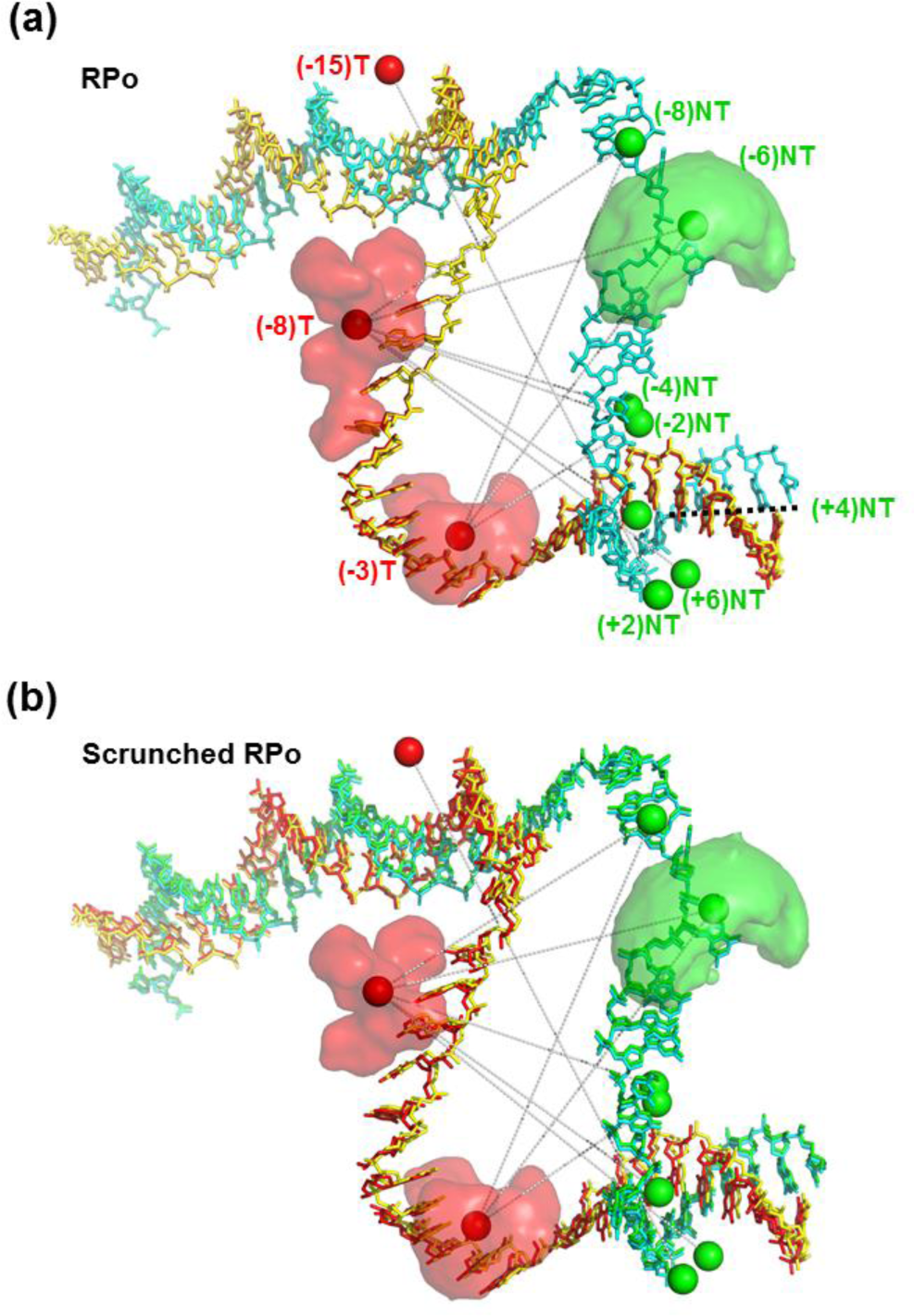
FPS analysis of transcription bubble conformational states. FPS analysis was performed for each conformational state against the known RPo structure (pdb code: 4XLN). (a) One set of apparent mean D-A distances, *r̅_D−A, E_* (*RPo state*) permits reconstruction of the template (T; red) and nontemplate (NT; green) strands in a 3D organization that matches the 3D organization of the T (yellow) and NT (cyan) strands in the crystal structure. (b), The second *r̅_D−A, E_* set (*scrunched RPo state*) yielded reconstruction of the template (T; red) and nontemplate (NT; green) strands in a 3D organization that does not match the 3D organization of the T (yellow) and NT (cyan) strands in the crystal structure. Solid spheres represent the mean positions of the D (green) and A (red) dyes. The set of mean D-A distances, *r̅_D−A, E_*, probed is shown by grey dashed lines connecting D and A mean positions. The mean position is always in within the dyes’ AVs (3 examples of dye AVs are shown).

## IV. Conclusions

### A. Transcription bubble dynamics and transcription start-site selection

Robb *et al.* have previously identified sub-millisecond transcription bubble dynamics using BVA^32^. In conjunction with their smFRET studies, they also performed radioactive gel-based transcription assays and have shown that transcription with the lacCONS promoter produced two bands of different-length transcripts. The longer transcript corresponded to the expected length assuming that transcription started from the canonical TSS. The ~ 2-3 bp shorter transcript suggested that bubble dynamics occurred between RPo and another conformation in which ~2-3 downstream DNA bp were scrunched, leading to a selection of a different transcription start site. We note, however, that transcription bubble dynamics and TSS selection could, in principle, occur independently. Our study, however, further corroborate Robb *et al.* conclusion: we showed that (i) the sub-millisecond dynamics is between two conformational states for the lacCONS promoter; (ii) one conformational state matches the known RPo crystal structure; (iii) the other conformational state is suggestive of DNA scrunching permitting the start of transcription at an alternative TSS, 2-3 bp downstream to the canonical TSS.

It is still unclear, however, whether the correlated increase in bubble coordinates and decrease in scrunching coordinates can occur without the melting of downstream DNA bases. Robb *et al.*’s study was followed by several studies that examined promoter sequence determinants for TSS selection^55–58^. These groups found specific promoter sequence characteristics that affect TSS selection. The mechanism suggested by Robb *et al.* and supported by our study may explain why TSS selection is a widespread phenomenon, with different types of scrunched RPo states, corresponding to different types of TSSs in different promoter sequences. Scrunching dynamics supporting TSS selection may depend also on the efficiency of the incorporation of the first NTP, from each RPo state. Our studies identified two conformational states, the *RPo state* and *scrunched RPo state* that are stable/survive for a few hundreds of microseconds. It is unclear, however, how efficient first NTP incorporation will be for short-lived conformations. To ultimately answer the question, an experiment that can correlate between TSS selection and bubble dynamics in real time is required.

### B. Determination of dynamic structures

In smFRET of immobilized molecules, state dwells, their FRET values and their transition rates could be directly retrieved from time trajectories but limited to rate constants <100 s^-1^ ^59, 60.^smFRET burst analysis of diffusing molecules can report on transition rate constants that are slower than the transit time through the observation volume (~1 ms), also effectively limited to ~100 s^-1^ ^20, 21^(Recent works have shown that faster dynamics, associated with rate constants up to ~5-10x10^3^ s^-1^, could be assessed using advanced technique ^21, 23, 24^ Nevertheless, since bright dyes can produce photon detection rates of ~1 MHz, photon-by-photon analysis could provide information on faster dynamics^25^. We note that interconversion between conformational states (separated by an activation barrier larger than k_B_T) could occur with even faster rate constants (~100x10^6^ s^-1^), but the amplitude of motion will be small at these very fast rates.

In this study we presented a strategy for the extraction of FRET-derived conformational states interconverting with rate constants of up to ~10x10^3^ s^-1^, that includes photon-by-photon analysis and relies on the utilization of a series of established tool ^17, 24, 25, 37.^ This strategy constitutes the following steps: (i) smFRET measurements are performed and the number of subpopulations in the FRET histogram is qualitatively identified. This step defines the minimal possible number of conformational states, *q*; (ii) BVA is applied to the data set to determine whether one (or more) of the FRET subpopulation(s) exhibit(s) FRET dynamics in within sm bursts. Presence of FRET dynamics hints the total number of states that may be larger than *q* (due to fast dynamics occurring while the molecule traverses the observation volume); (iii) photon-by-photon hidden Markov modelling (H2MM) analysis is applied to photon bursts with models of increasing number of states (starting from the q-state model). H2MM analyses results are assessed by comparing the values of a modified Bayes information criterion (*BIC’;* see Eq. S11 of the supplementary material) to determine the minimal sufficient number of states; (iv) the mean FRET efficiency values ; are extracted for the H2MM identified states (Eq. S16 of the supplementary material) and transformed into apparent mean D-A distances *r̅_D−A, E_* (Eqs. S17, S18, S21 of the supplementary material); (v) steps (i)-(iv) are repeated for all labeled constructs; The more labeled constructs of more reaction coordinates are measured, the more accurate and reliable the dynamic structure determination will be; (vi) the different apparent mean D-A distances of different labeled constructs are classified and grouped to represent different conformational states; (vii) grouped *r̅_D−A, E_* value sets are used as constraints in FPS to determine dynamic conformations.

In cases where crystal structures exist, FPS-determined conformations can be compared to the crystal structures and identified. FPS conformations that do not match the crystal structure might be destabilized in the crystallization process. Even when crystal structures of specific conformations do not exist, the same procedure could be used against structures computed by molecular dynamics (MD)^19, 43–48,^ Monte Carlo (MC)^19, 49^or coarse-grained (CG)^49–51^ simulations or against database predicted structures^52^.

A standardization effort for smFRET-based structure determination, led by the Hügel and Seidel groups, was initiated following the wwPDB Hybrid/Integrative methods task force recommendations^61^. The approach presented here extends the above-mentioned efforts and outlines a general approach for assessing rapidly interconverting conformations. We demonstrated the approach for the transcription bubble at RPo with the lacCONS promoter. Due to its generality it could be used to elucidate interconverting conformations of other macromolecules.

## V. SUPPLEMENTARY MATERIAL

The supplementary material include description of the μsALEX-smFRET approach (supplementary text, section A), description of the BVA method (supplementary text, section B), description of the H2MM approach (supplementary text, section C), description of most likely *PR* to mean FRET efficiency conversion protocol (supplementary text, section D), description of FRET efficiencies to D-A distances conversion protocol (supplementary text, section E), description of the protocol for diffusion based smFRET simulations (supplementary text, section F), steady-state fluorescence measurements results for singly-labeled (D or A) lacCONS promoter (FIG. S1), BVA results (FIG. S2), promoter opening and escape activity results (FIG. S4), examples of constructs that failed the screening criteria (FIGs. S3, S5-S7), H2MM state-model assessment example (FIG. S8, S9), diffusion-based simulation (FIG. S11) and their results (FIG. S10), analyses of control measurements (FIGs. S12-S14), an example of dwell analysis on Viterbi-produced trajectories (FIG. S15), assessment of the effect of BG correction on mean *PR* values (FIG. S16), assessment of the inaccuracy in using the Förster relation (Eq. S17) to directly calculate the mean D-A distance from mean FRET efficiencies (FIG. S17), correction factors (Table S1), H2MM analysis results of main experiments (Table S2), diffusion-based smFRET simulated data (Table S3) and of control measurements (Tables S4, S5).

In this work we analyzed sm bursts, performed BVA and generated FIGs. 2, S2, S4 and S10 using the FRETBursts software^62^. FRETBursts Python Notebooks that include the analyses and generation of these figures are deposited in Figshare^63–66^, The raw data used to generate these figures was saved in Photon-HDF5 format^67^ and deposited in Figshare as well^68–70^. Using the ALiX software^71^, we extracted the photon stream identity and timestamp information, for all photons in the selected bursts^72, 73.^These files were used as the base input for the H2MM analysis. The results of the H2MM analyses are stored in Matlab files^74, 75.^In addition, the core code for the H2MM analysis was provided by Pirchi, Tsukanov *et al.^25^* Additional matlab scripts for dwell analysis (including additional Viterbi algorithm code, generously provided by Dr. Menachem Pirchi) are also deposited in Figshare, including code documentation and script usage instructions^76^. Brownian motion-based simulations of smFRET experiments including 2-state dynamics were performed with the software PyBroMo^77^. All pdb files used in FPS ^78, 79^,input data^80^, analysis results for *RPo^81^* and *scrunched RPo^82^* states, and PyMOL scripts used for preparation of FIG. 4 ^83^, are also stored in Figshare.

## VI. ACKNOWLEDGEMENTS

We would like to thank Dr. Xavier Michalet, Dr. SangYoon Chung, Mr. Yazan Alhadid, Prof. Arieh Warshel and Dr. Raphael Alhadeff for fruitful discussions, Dr. Xavier Michalet for consultation regarding ALiX software and Dr. Menachem Pirchi for consultation regarding H2MM. We thank Prof. Thorben Cordes for generously providing us with some of the labeled DNA oligos. This work was supported by the NIH under grant numbers GM069709 (to SW) and GM095904 (to XM and SW); NSF under grant number MCB-1244175 (to SW).

